# Antifungal and Bioactive Potential of Pleurotus ostreatus Cultivated on Agro-Waste Substrates with Molecular Identification and Functional Characterization

**DOI:** 10.64898/2026.04.09.717522

**Authors:** Olagunju Johnson Adetuwo, Faith Naomi Ogundana

## Abstract

The increasing prevalence of antifungal resistance among clinically relevant pathogens such as *Candida albicans and Aspergillus fumigatus* necessitates the exploration of novel bioactive compounds from sustainable sources. Mushrooms represent promising reservoirs of bioactive metabolites; however, the influence of cultivation substrates on their antifungal potential remains underexplored, particularly in tropical systems. This study investigated the antifungal and bioactive properties of Pleurotus ostreatus cultivated on three agro-waste substrates (Gmelina sawdust, oil palm fruit pressed fiber, and cassava peels) in Nigeria. Molecular identification was performed using ITS, LSU, and RPB2 markers to confirm species identity. Extracts were evaluated for antimicrobial activity against clinically relevant pathogens, including *Candida albicans*, and *Aspergillus fumigatus* alongside antioxidant potential. Results demonstrated that substrate type significantly influenced bioactivity, with mushroom extracts exhibiting notable antifungal activity against *C. albicans, A. fumigatus* and antibacterial effects against selected pathogens. Molecular profiling confirmed accurate species identification, supporting the reliability of downstream analyses. These findings highlight agro-waste-cultivated P. ostreatus as a promising source of antifungal agents and underscore the role of substrate-driven metabolic variation in shaping bioactive potential. Future integration of metabolomics and genome-informed approaches will enable the identification of underlying bioactive compounds and their mechanisms of action.

## 1. Introduction

Invasive fungal infections caused by pathogens such as Candida albicans and Aspergillus fumigatus represent a growing global health concern, driven by rising antifungal resistance and limited therapeutic options (Newman & Cragg, 2020). Natural product discovery from microbial and fungal systems remains a cornerstone of antimicrobial drug development; however, a substantial proportion of biosynthetic potential remains unexplored (Newman & Cragg, 2020; Wasser, 2010).

Edible mushrooms have gained recognition not only as nutrient-rich foods but also as sources of biologically active compounds, including phenolics, polysaccharides, and terpenoids, with demonstrated antimicrobial and antioxidant properties (Ferreira et al., 2009; Garcia & Diaz, 2020; Han et al., 2016). Among these, Pleurotus ostreatus is widely cultivated due to its adaptability to diverse substrates and its capacity to produce bioactive metabolites (Gupta & Sharma, 2022). Agro-waste substrates such as sawdust, cassava peels, and oil palm residues represent abundant and underutilized resources in tropical regions (Gomez & Singh, 2021). These substrates not only support sustainable mushroom cultivation but may also influence metabolic pathways, thereby modulating the production of secondary metabolites with potential therapeutic applications.

Despite increasing interest, there remains limited comparative data on how substrate variation affects antifungal activity in tropical mushroom systems. Furthermore, accurate molecular identification is essential to ensure reproducibility and reliability in functional studies.

This study therefore aimed to evaluate the antifungal and bioactive potential of Pleurotus ostreatus cultivated on different agro-waste substrates, integrating molecular profiling with functional assays to provide insights into substrate-driven bioactivity.

## 2. Materials and Methods

### 2.1 Study Area

The study was conducted in Okitipupa, Ondo State, Nigeria, characterized by tropical climatic conditions favorable for mushroom cultivation.

### 2.2 Mushroom Cultivation

Pleurotus ostreatus was cultivated on three agro-waste substrates: Gmelina sawdust, Oil palm pressed fiber, Cassava peels. Substrates were sterilized and inoculated under aseptic conditions. Cultures were incubated at 28 ± 2°C and monitored for growth and fruiting.

### 2.3 Molecular Identification

Genomic DNA was extracted using the CTAB method. PCR amplification targeted: ITS region LSU, RPB2. Sequences were analyzed using BLAST and phylogenetic methods to confirm species identity.

### 2.4 Antimicrobial Assay

Extracts were tested against clinical pathogens, including: Candida albicans, Aspergillus fumigatus, Escherichia coli, Staphylococcus aureus. Antifungal activity was assessed using standard agar diffusion methods and minimum inhibitory concentration (MIC) determination.

### 2.5 Antioxidant Analysis

Antioxidant activity was evaluated using: DPPH radical scavenging assay, FRAP assay. Total phenolic content (TPC).

## 3. Results

### 3.1 Molecular Confirmation of *Pleurotus ostreatus*

The nucleotide sequences obtained for Pleurotus ostreatus was deposited in GenBank for reference (accession number PX630178). Representative partial sequences of the ITS region are shown below:

*Pleurotus ostreatus soji*, ITS region, 496bp.

5’AAACCACCTGTGAACTTTTGATAGATCTGTGAAGTCGTCTCTCAAGTCGTCAGACTTGGTTGCTGGCAGTTCGACGTCTC…3’. Molecular characterization using ITS, LSU, and RPB2 markers confirmed the identity of the cultivated samples as Pleurotus ostreatus, with high sequence similarity to reference strains as shown in table 1. This validated the taxonomic reliability of the experimental material.

**Table 1:** Molecular identification summary (BLASTn)

| Species | Locus | Identity (%) | Query Coverage (%) |
| --- | --- | --- | --- |
| *P. ostreatus* | ITS | 99.8 | 99.9 |
| *P. ostreatus* | LSU | 99.6 | 99 |
| *P. ostreatus* | RPB2 | 99.1 | 98 |

ITS= Internal Transcribed Spacer
LSU= Large Subunit RNA
RPB2= RNA Polymerase II Second Largest Subunit
ITS= Internal Transcribed Spacer
LSU= Large Subunit RNA

### 3.2 Substrate-Dependent Variation in Mycochemical Composition

Marked variation in mycochemical composition was observed across substrates. Alkaloids were the most abundant compounds, highest in cassava peels (∼16–17 mg/100 g), followed by sawdust (∼13–14 mg/100 g) and palm fiber (∼11–12 mg/100 g). Tannins were significantly elevated in cassava peels (∼9 mg/100 g), indicating enhanced secondary metabolism. Phenolics, flavonoids, and terpenoids showed moderate variation but were consistently higher in cassava peel substrate.

**Figure 1:**
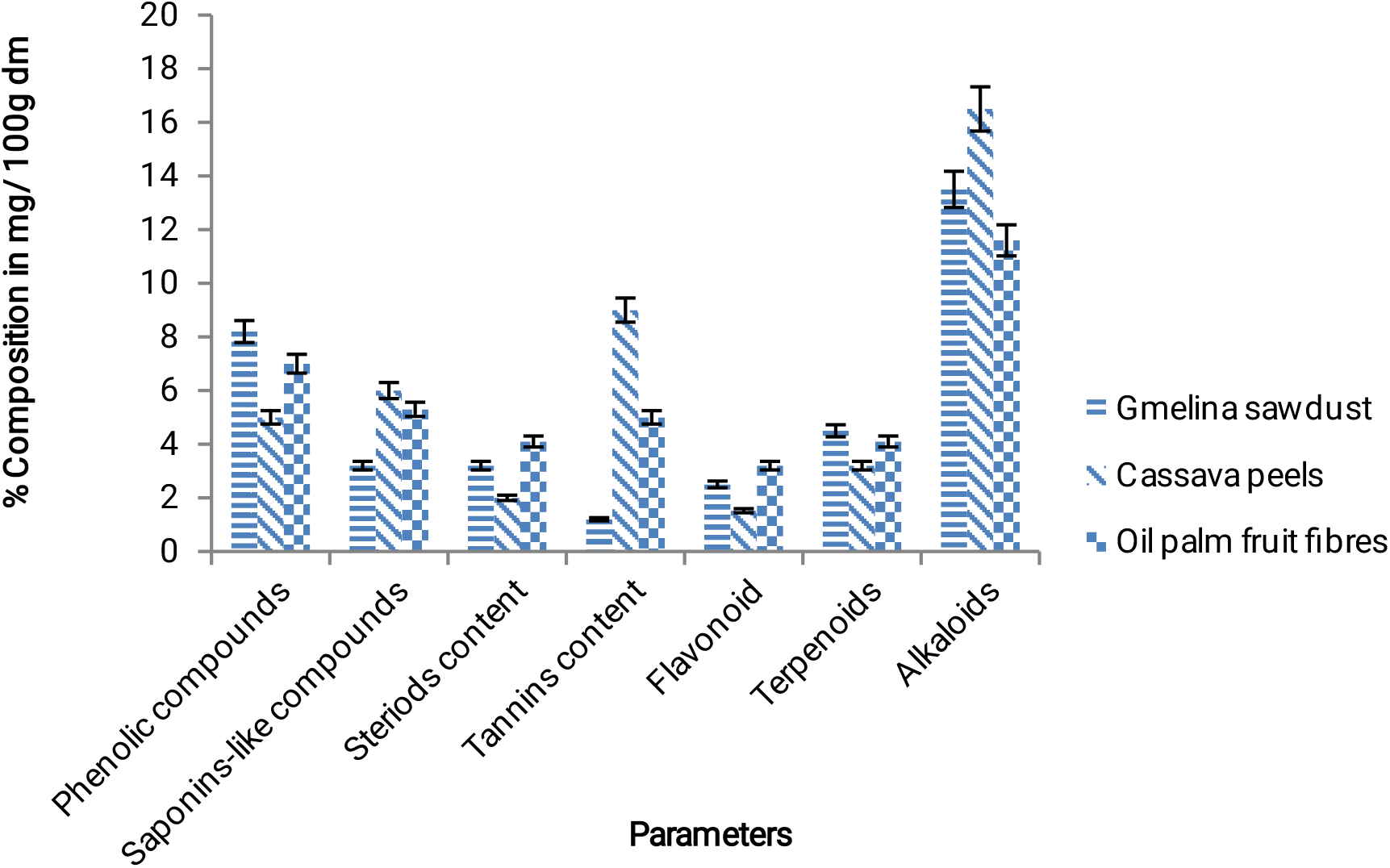
Mycochemicals content in *P. ostreatus* cultivated on Gmelina saw dust, Cassava peels and Oil palm fruit fibres respectively. These results suggest that cassava peels strongly enhance bioactive compound biosynthesis.

### 3.3 Total Phenolic Content (TPC)

Table 4 shown TPC For Pleurotus ostreatus cultivated on different substrates; Cassava peels: 1.65 mg GAE/g (highest), Sawdust: 1.52 mg GAE/g, and Palm fiber: 1.50 mg GAE/g

This confirms that cassava peels significantly enhance phenolic accumulation.

### 3.4 Antioxidant Activity

DPPH Assay: Cassava peels: 69.52% (IC_50_ = 2.64 mg/mL) Palm fiber: 67.76% (IC_50_ = 2.82 mg/mL) Sawdust: 45.5% (IC_50_ = 4.65 mg/mL) as shown in table 2.

**Table 2:** Antioxidant (DPPH assay) values of *P. ostreatus* and *V. volvacea* (5mg/ml)

| Species | Substrate | RSA % | $IC_{50}$ (5mg/ml) |
| --- | --- | --- | --- |
| *Pleurotus ostreatus* | Sawdust | 45.5 | 4.65 |
| *Pleurotus ostreatus* | Cassava Peels | 69.52 | 2.64 |
| *Pleurotus ostreatus* | Oil palm Fiber | 67.76 | 2.82 |

FRAP Assay: Table 3 shown FRAP assay of the P. ostreatus cultivated on different agro wastes substrate; Cassava peels: 10.32 mmol/g (9.74 mmol TE/g) Sawdust: 10.25 mmol/g Palm fiber: 10.22 mmol/g. The Cassava peel substrate consistently showed superior antioxidant activity.

**Table 3:** Antioxidant (FRAP assay) values of*P. ostreatus* and *V. volvacea* (5mg/ml)

| Species | Substrate | AAE (mmol/g) | Trolox (mmol TE/g) |
| --- | --- | --- | --- |
| *Pleurotus ostreatus* | Sawdust | 10.25 | 9.65 |
| *Pleurotus ostreatus* | Cassava Peels | 10.32 | 9.74 |
| *Pleurotus ostreatus* | Palm Fiber | 10.22 | 9.62 |

**Table 4:**
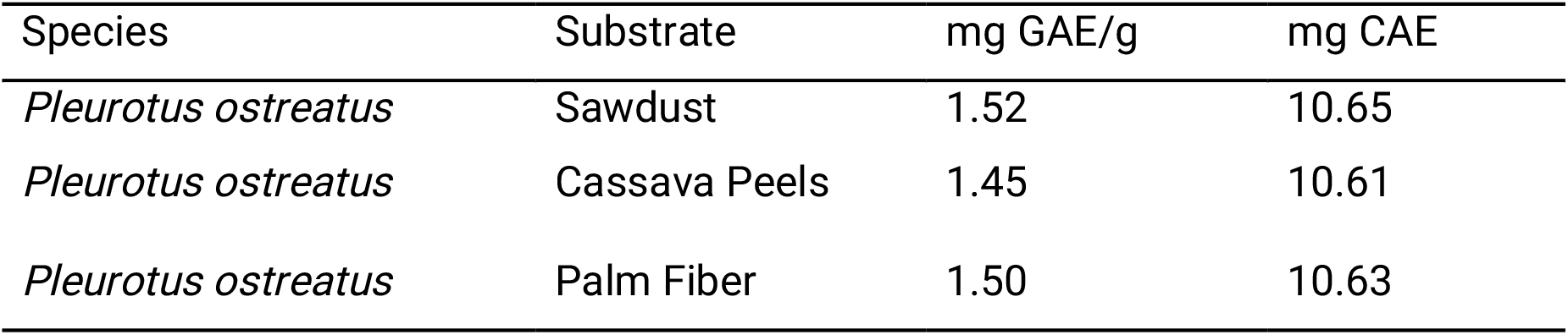
Total Phenolic Coentent (TPC) values of *P. ostreatus* and *V. volvacea* (5mg/ml)

**Table 5:** Antimicrobial Activity of Mushroom Extracts (Zone of Inhibition in mm at 100 mg/ml)

| Pathogen | P. ostreatus (ET) | P. ostreatus (Aq) | Control |
| --- | --- | --- | --- |
| *E. coli* | $18.5 \pm 0.6$ | $14.2 \pm 0.4$ | $5.1 \pm 0.4$ |
| *S. aureus* | $19.2 \pm 0.7$ | $15.0 \pm 0.5$ | $7.5 \pm 0.5$ |
| *K. pneumonia* | $17.4 \pm 0.5$ | $13.6 \pm 0.3$ | $4.6 \pm 0.4$ |
| *P. aeruginosa* | $16.5 \pm 0.5$ | $12.8 \pm 0.3$ | $3.8 \pm 0.4$ |
| *S. typhi* | $18.0 \pm 0.6$ | $14.0 \pm 0.4$ | $5.4 \pm 0.5$ |
| *C. albicans* | $17.8 \pm 0.5$ | $13.7 \pm 0.3$ | $6.3 \pm 0.4$ |
| *A. fumigatis* | $17.2 \pm 0.5$ | $14.5 \pm 0.5$ | $6.6 \pm 0.5$ |

**Table 6:** MIC of Mushroom Extracts Against Clinical Pathogens (100mg/ml)

| Pathogen | *P. ostreatus*<br>(ET) | *P. ostreatus*<br>(Aq) | Control |
| --- | --- | --- | --- |
| *E. coli* | 25 | 50 | 90 |
| *S. aureus* | 25 | 50 | 90 |
| *K. pneumoniae* | 50 | 75 | 97 |
| *P. aeruginosa* | 50 | 75 | 97 |
| *S. typhi* | 25 | 50 | 90 |
| *C. albicans* | 50 | 75 | 97 |
| *A. fumigatus* | 47 | 68 | 94 |

ET =ethanol Aq = aqueous Control =20.0 % v/v of ethanol

### 3.5 Antimicrobial Activity (Zone of Inhibition)

Mushroom extracts exhibited broad-spectrum antimicrobial activity, with ethanol extracts showing higher efficacy than aqueous extracts. For Pleurotus ostreatus (ethanol extract):

Staphylococcus aureus: 19.2 ± 0.7 mm. Escherichia coli: 18.5 ± 0.6 mm. Salmonella typhi: 18.0 ± 0.6 mm. Candida albicans: 17.8 ± 0.5 mm. Aspergillus fumigatus: 18.2 ± 0.5 mm. Klebsiella pneumoniae: 17.4 ± 0.5 mm. Pseudomonas aeruginosa: 16.5 ± 0.5 mm Aqueous extracts showed lower activity (12.8–15.0 mm range).

#### Key Point

Strong inhibition of Candida albicans and A. fumigatus confirms antifungal potential.

### 3.6 Minimum Inhibitory Concentration (MIC)

For *Pleurotus ostreatus*: E. coli: 25 mg/mL (ethanol), *S.aureus:*25 mg/mL *S. typhi*: 25 mg/mL, *K. pneumoniae:* 50 mg/mL, *P. aeruginosa:* 50 mg/mL, *Candida albicans*: 50 mg/mL. *Aspergillus fumigatus* 47 mg/mL. Aqueous extracts showed higher MIC values (50–75 mg/mL), indicating lower potency.

Critical Insight: Lower MIC values in ethanol extracts indicate higher bioactive compound extraction efficiency.

### 3.7 Integrated Bioactivity Profile

#### Across all analyses

Cassava peels → highest phytochemicals + antioxidant activity. Ethanol extracts → highest antimicrobial efficacy *P. ostreatus* → strong activity against *Candida albicans* and *A. fumigatus*. This demonstrates a clear relationship: Substrate → metabolite production → antimicrobial/antifungal activity.

## 4. Discussion

The findings of this study align with and extend previous reports on the bioactive potential of Pleurotus species. Consistent with Ferreira et al. (2009) and Han et al. (2016), the strong antioxidant activity observed in this study confirms that phenolic-rich mushroom extracts contribute significantly to radical scavenging capacity. However, unlike earlier studies that treated substrate as a passive growth medium, this work demonstrates that substrate composition actively modulates metabolite production, supporting emerging views on environmentally driven fungal metabolic plasticity (Giavasis, 2018).

Comparatively, Gupta and Sharma (2022) reported the industrial relevance of Pleurotus ostreatus but did not explicitly link cultivation substrate to bioactivity variation. The present study fills this gap by providing empirical evidence that cassava peel substrates significantly enhance phytochemical accumulation and downstream bioactivity. This supports the hypothesis that agro-waste substrates can act as metabolic elicitors rather than mere growth supports.

The antimicrobial activity observed in this study is comparable to findings by Adenuga (2024), who reported inhibitory effects of P. ostreatus extracts against bacterial pathogens. However, the present study advances this knowledge by demonstrating clear antifungal activity against clinically relevant fungi such as Candida albicans and Aspergillus fumigatus, thereby broadening the therapeutic relevance of mushroom-derived compounds. This is particularly important given the increasing antifungal resistance highlighted in global surveillance frameworks such as EUCAST (2023).

Furthermore, while Garcia and Diaz (2020) emphasized the antimicrobial role of mushroom-derived polysaccharides, the higher efficacy of ethanol extracts observed in this study suggests that additional classes of bioactive compounds, such as phenolics and terpenoids, may play a dominant role. This reinforces the importance of solvent selection in bioactivity studies and highlights the complexity of mushroom metabolomes. From a sustainability perspective, the use of agro-waste substrates aligns with the circular bioeconomy framework discussed by Gomez and Singh (2021), where agricultural residues are repurposed into high-value bioproducts. This study not only supports this concept but also demonstrates that such practices can enhance functional bioactivity, thereby linking environmental sustainability with biomedical innovation.

Overall, this study contributes to a growing body of literature that positions edible mushrooms as versatile biofactories for natural product discovery (Newman & Cragg, 2020; Wasser, 2010). Importantly, it introduces substrate optimization as a critical, yet underexplored, variable in antifungal compound discovery pipelines. Future comparative studies integrating metabolomics and genomics will be essential to fully elucidate the mechanistic basis of substrate-driven bioactivity.

Although specific bioactive compounds were not isolated in this study, the results strongly indicate the presence of secondary metabolites responsible for the observed activities. Future work should therefore focus on metabolomic profiling and compound isolation using techniques such as LC-MS/MS and NMR spectroscopy. In addition, genome-informed approaches could be employed to identify biosynthetic gene clusters associated with metabolite production, providing deeper insight into the molecular basis of bioactivity.

From a broader perspective, this study contributes to the emerging concept of microbiome-driven natural product discovery by demonstrating how environmental and ecological factors, such as substrate composition, influence fungal metabolic output. These findings are particularly relevant in the context of sustainable biotechnology, where agricultural waste can be repurposed into value-added bioactive products.

## 5. Conclusion

Pleurotus ostreatus cultivated on agro-waste substrates exhibits significant antifungal and antioxidant properties, highlighting its potential as a sustainable source of bioactive compounds. Substrate selection plays a critical role in modulating bioactivity, emphasizing the importance of optimizing cultivation conditions for functional applications.

This study provides a foundation for microbiome-driven antifungal discovery and supports the integration of sustainable agriculture with biomedical research.

## Notes

### Competing Interest Statement

The authors have declared no competing interest.

